# The role of rice SOG1 and SOG1-like in DNA damage response

**DOI:** 10.1101/2022.01.21.477278

**Authors:** Ayako Nishizawa-Yokoi, Ritsuko Motoyama, Tsuyoshi Tanaka, Akiko Mori, Keiko Iida, Seiichi Toki

## Abstract

Higher plants are constantly exposed to environmental stresses, and therefore complicated defense systems, including DNA damage response (DDR) and DNA repair systems, have developed to protect plant cells. In *Arabidopsis*, the transcription factor SUPPRESSOR OF GAMMA RESPONSE 1 (SOG1) has been reported to play a key role in DDR. Here, we focus on DDR in rice—thought to be a simpler system compared with *Arabidopsis* due to lack of induction of endocycle even under DNA damage stress. Rice SOG1 (OsSOG1) and SOG1-like (OsSGL) were identified as putative AtSOG1 orthologs with complete or partial conservation of the serine-glutamine (SQ) motifs involved in activation via phosphorylation. In addition to OsSOG1- or OsSGL-knockout mutants, OsSOG1 non-phosphorylatable mutants (OsSOG1-7A) were generated by homologous recombination-mediated gene targeting. Based on DNA damage susceptibility and transcriptome analysis using these mutants, we demonstrated that OsSOG1, but not OsSGL, plays a central role in the DDR and DNA repair. OsSOG1 regulated target genes via CTT (N)_7_ AAG motifs reported previously as AtSOG1 recognition sites. The loss of transcription activities and DNA damage tolerance of OsSOG1-7A was not complete compared with OsSOG1-knockout mutants, raising the possibility that another phosphorylation site might be involved in the activation of OsSOG1. Furthermore, our findings have highlighted differences in SOG1-mediated DDR between rice and *Arabidopsis*, especially regarding induction of cell-cycle arrest and endocycle arrest, revealing rice-specific DDR mechanisms.

**One sentence summary:** Rice transcription factor SUPPRESSOR OF GAMMA RESPONSE 1 controls DNA damage response and DNA repair through activation via phosphorylation and the direct regulation of expression of numerous genes.

## Introduction

The sessile nature of higher plants results in constant exposure to environmental stresses; therefore, complex defense systems and/or signaling networks have been developed to protect cells from damage in response to stress. DNA damage response (DDR) and repair systems are among the most important defenses against environmental stresses such as ultraviolet B irradiation, drought, salt, heat, and other pollutants, e.g., metals. Reactive oxygen species (ROS) generated in response to environmental stresses alter bases and damage sugar residues, resulting in DNA fragmentation and strand breaks (Roldán-Arjona and Ariza, 2009). DNA double-strand breaks (DSBs) are the most severe type of damage because they lead to cell death if not repaired. DSBs are repaired by two distinct pathways: non-homologous end-joining (NHEJ), which simply ligates DNA ends, and homologous recombination (HR), which repairs DSBs precisely by copying a donor DNA molecule. The mechanisms of DDR have been well studied in mammalian cells. According to previous studies, the tumor suppressor protein p53—the “guardian of the genome”—plays a pivotal role in a variety of DDRs, including the promotion of cell cycle arrest, the induction of apoptosis, and regulating DNA repair (Ventura et al., 2007; Reinhardt and Schumacher, 2012). However, p53 orthologues are not conserved in plants (Yoshiyama et al., 2013), and DDR in plant cells remains poorly understood, except in *Arabidopsis.*

In *Arabidopsis*, SUPPRESSOR OF GAMMA RESPONSE 1 (SOG1), an NAC (No apical meristem, Arabidopsis transcription activation factor, Cup-shaped cotyledon) transcription factor, has been reported to regulate the expression of numerous genes relating to DNA repair, cell cycle arrest and stem cell death under DNA damage conditions. Thus, SOG1 is thought to play a key role in DDR and to be a counterpart of mammalian p53 (Yoshiyama et al., 2009, 2013; Ogita et al., 2018). Similarly to p53, *Arabidopsis* SOG1 (AtSOG1) is activated via phosphorylation of a serine-glutamine (SQ) motif in the C-terminal region of AtSOG1 by the cellular kinases ATM (ATAXIA-TELANGIECTASIA MUTATED) and ATR (ATM AND RAD3-RELATED), which are involved in the sensing of DNA lesions upon treatment with DSB-inducing agents (Yoshiyama et al., 2013, 2017; Sjogren et al., 2015; Yoshiyama and Kimura, 2018). Predicted SOG1 ortholog proteins with conserved SQ motif phosphorylated by ATM and ATR are found in most land plants, from moss to angiosperm (Yoshiyama et al., 2014). Wei et al. (2021) reported recently that *Arabidopsis* casein kinase (CK) 2 modulated AtSOG1 activity through its phosphorylation under DNA damage conditions, and CK2 phosphorylation sites ([ST]XX[ED]) are conserved among putatively orthologous proteins in several plant species (Wei et al., 2021). Moreover, the same study also demonstrated that inhibition of CK2-dependent phosphorylation of AtSOG1 interfered with its phosphorylation by ATM/ATR following DNA damage (Wei et al., 2021). Several recent studies have reported that SOG1 is also involved in DDR derived from oxidative stress including aluminium- (Sjogren et al., 2015) and cadmium- (Hendrix et al., 2020) exposure or biotic stress (Ogita et al., 2018) in *Arabidopsis.* These findings suggest that *Arabidopsis* SOG1 regulates several biological processes in response to the DNA damage that occurs under various environmental stress conditions via its phosphorylation by ATM/ATR or CK. However, there have been hardly any studies on either the transcriptional regulatory or activation mechanisms via phosphorylation of SOG1 in response to DNA damage in plant species other than *Arabidopsis*.

Even though rice is one of the most agronomically important crops, the mechanisms of DDR in rice remain to be understood. Unlike in *Arabidopsis*, DDR in rice is thought to be simple, because the process of endocycle is not found in rice other than during endosperm development, even under stress conditions (Endo et al., 2012). Thus, it is expected that rice can safeguard genome integrity against environmental stresses by the repair of damaged DNA or by the induction of cell death. These differences raise interesting questions about DDR and the function of SOG1 in rice. In addition, genome editing techniques, not only clustered regularly interspaced short palindromic repeats (CRISPR)/CRISPR-associated 9 (Cas9)-mediated targeted mutagenesis for producing loss-of-function mutants by the induction of insertion or deletion mutations at CRISPR target site, but also HR-mediated gene targeting (GT) for producing gain-of-function mutants by the introduction of point mutations or the insertion of reporter gene at the desired locus, are now well-developed and make it possible to analyze precisely a characteristic feature of a target gene in rice.

We have identified rice SOG1 (OsSOG1, Os06g0267500), which carries the five highly conserved SQ motifs of AtSOG1, and SOG1-like (OsSGL, Os02g0594800), which partially retains SQ (or threonine-glutamine, TQ) motifs from the rice genome, by a homology search of public databases (**Supplementary Figure 1**). In *Arabidopsis*, ANAC044 and ANAC085 are identified as the closest homologs of AtSOG1, but lack SQ motifs (Takahashi et al., 2019). A total of 151 NAC genes are predicted to be present in the rice genome (Nuruzzaman et al., 2010). OsSOG1 and OsSGL were designated as ONAC091 and ONAC050, respectively, in the Rice Annotation Project (RAP) database (https://rapdb.dna.affrc.go.jp/index.html). However, no previous functional analyses of these transcription factors have been reported. We generated OsSOG1- and OsSGL-knockout rice plants by GT and also produced OsSOG1/OsSGL double knockout mutants by CRISPR/Cas9-mediated targeted mutagenesis. In addition to these knockout mutants, non-phosphorylatable and phosphomimetic OsSOG1 mutants substituting the serine residues for alanine or aspartic acid, respectively, at all SQ motifs of *OsSOG1* gene were generated by GT to investigate the importance of the SQ motifs of OsSOG1 in DDR. These mutants, generated via GT, harbored a modified OsSOG1 gene with substitutions of phosphorylation sites at its endogenous locus, thereby allowing the activation mechanisms via phosphorylation of OsSOG1 in response to DNA damage to be assessed precisely without being affected by position effect variegation.

In this report, we evaluated DNA damage susceptibility and transcriptome analysis using a single knockout mutant of OsSOG1 or OsSGL, a double knockout mutant of OsSOG1 and OsSGL, and a non-phosphorylatable mutant of OsSOG1. Unfortunately, we were unable to obtain OsSOG1 phosphomimetic mutants by GT. OsSOG1 was demonstrated to be a pivotal factor regulating the transcription of genes involved in DNA repair and cell cycle processes through a specific motif reported previously as the SOG1-binding domain in *Arabidopsis.* Based on these results, we report that OsSOG1, but not OsSGL, plays a central role not only in DDR and DNA repair but also in the oxidative stress response in rice. Although SQ motifs of OsSOG1 were important in the function of OsSOG1 in DDR, our findings suggest the existence of another pathway (e.g., via CK) in the activation of OsSOG1 during DNA damage.

In addition, our findings suggest that monocotyledonous rice might overcome DNA damage using different mechanisms from dicotyledonous *Arabidopsis*, e.g. cell-cycle arrest. Knowledge gained from our studies could help elucidate mechanisms of DDR in rice—one of our most important agricultural crops—and has significant implications with respect to the development of genome editing techniques that exploit DNA repair pathways for crops.

## Results

### Generation of OsSOG1 and OsSGL mutant rice by HR-mediated GT and/or CRISPR/Cas9-mediated mutagenesis

To investigate the function of OsSOG1 and OsSOG1-like (OsSGL) in DDR and DNA repair, we generated several types of OsSOG1 and OsSGL mutant rice plants. OsSOG1 promoter::β-glucuronidase (Psog1-GUS) and OsSGL promoter::GUS (Psgl-GUS) knock-in (simultaneous knock-out of OsSOG1 and OsSGL, respectively) rice mutants were generated by the introduction of a GUS::terminator construct in-frame with the ATG codon of *OsSOG1* or *OsSGL* by HR-mediated GT with positive-negative selection (Terada et al., 2002; Nishizawa-Yokoi et al., 2015) (**Figure 1A and Figure S2A**). Two independent lines each of Psog1-GUS (#1 and #2) and Psgl-GUS (#1 and #2) were obtained, and their homozygous mutant plants in the T_2_ or T_3_ generation were subjected to further analysis (**Figures 1 B, C and S2B, C**). No full-length transcripts of OsSOG1 and OsSGL were detected in Psog1-GUS and Psgl-GUS homozygous mutant plants, respectively, by reverse transcriptase-PCR (RT-PCR) (**Figure S3**).

**Figure 1.**
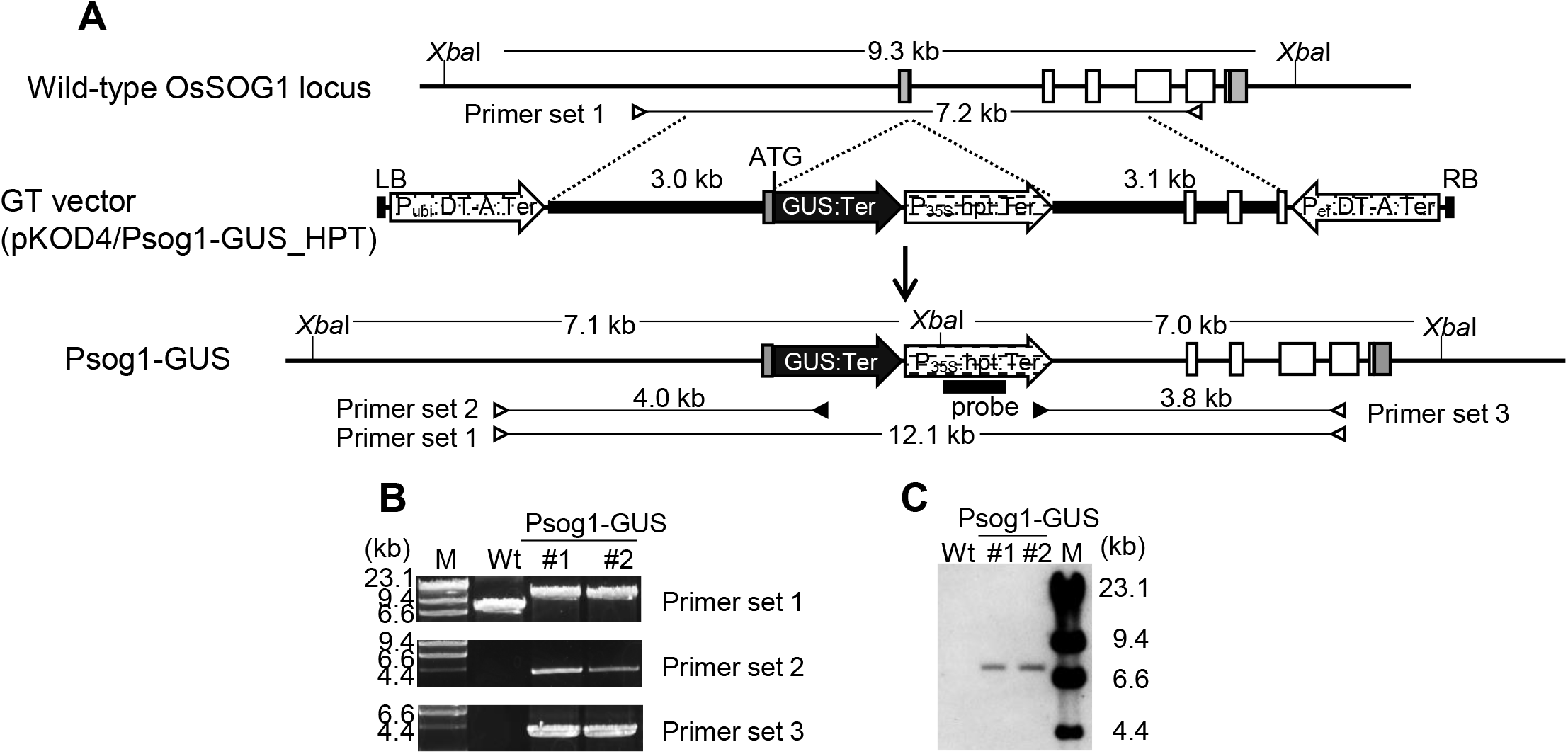
The generation of Psog1-GUS mutants by knock-in of the GUS gene downstream of the OsSOG1 promoter via GT. A, Strategy for the knock-in of a GUS::terminator construct in-frame with the translational start codon (ATG) of the *OsSOG1* gene by GT with positive-negative selection. The top line indicates the genomic structure of the wild-type *OsSOG1* gene region. Gray and white boxes show untranslated regions and exons, respectively. The middle line shows a schematic of the GT vector carrying two expression cassettes of diphtheria toxin A subunit gene (*DT-A*) under the control of the maize polyubiquitin 1 promoter (P_ubi_) or rice elongation factor 1α promoter (P_ef_) located on both ends as a negative selection marker, a GUS gene::terminator cassette, and the cauliflower mosaic virus 35S promoter (P_35S_)::hpt::terminator expression cassette as a positive selection marker within the sequence homologous to the *OsSOG1* locus (promoter region, 3.0 kb; coding region, 3.1 kb). LB, left border; RB, right border. The bottom line represents the Psog1-GUS mutant locus resulting from HR between the GT vector and wild-type locus. Primer sets 1, 2, and 3 were used for PCR analysis to identify the Psog1-GT mutants. *XbaI* was the restriction enzyme used for Southern blot analysis. B, PCR analysis with wild-type and two independent lines of Psog1-GUS T_2_ homozygous mutants (#1 and #2) using primer sets 1, 2, and 3 shown in A. C, Southern blot analysis with the hpt-specific probe shown in A using genomic DNA digested with *Xba*I.

A previous study demonstrated that ATM-mediated phosphorylation of the SQ motif in the C-terminal region of AtSOG1 plays a pivotal role in DDR in Arabidopsis (Yoshiyama et al., 2013). Yoshiyama et al. (2017) also revealed that the extent of SQ phosphorylation in SOG1 regulates gene expression levels and determines the strength of DDR (Yoshiyama et al., 2017). Therefore, we substituted the serine residues with alanine (non-phosphorylatable mutations) at all seven SQ motifs of the *OsSOG1* gene via GT with positive-negative selection and subsequent marker-excision by *piggyBac* transposition (Nishizawa-Yokoi et al., 2015) (**Figure 2A**). These mutant lines were called mutants OsSOG1–7A. After marker excision, seven amino acid substitutions (S360A, S316A, S322A, S338A, S340A, S398A, and S403A) and a silent mutation for the production of a *HpaI* restriction enzyme recognition site used for the insertion of *piggyBac* transposon with the positive selection marker were left in the *OsSOG1* gene locus (**Figure 2B**). The *piggyBac* transposase expression cassette was transformed into a GT callus line for marker excision from the *OsSOG1* locus; thus, T_1_ progeny plants with the desired mutations in the *OsSOG1* locus in the homozygous state and without transgene were selected by PCR. Their homozygous mutant progenies in the T_2_ or T_3_ generation were subjected to further analysis (**Figure 2C**). RT-PCR revealed transcript levels of the *OsSOG1* gene in OsSOG1-7A mutants to be comparable with those of wild-type (**Figure S3**). We also attempted to introduce substitution of the serine residues to aspartic acid (phosphomimetic mutations) at all seven SQ motifs of OsSOG1 by CRISPR/Cas9-mediated GT in rice (Nishizawa-Yokoi et al., 2020). Hygromycin-resistant callus lines propagated clonally from calli transformed with GT vector were subjected to cleaved amplified polymorphic sequence (CAPS) screening for isolation of GT calli (**Figure S4A**), since the additional silent mutation was introduced together with amino acid substitutions into the 5th exon of *OsSOG1* via GT to produce a *PstI* restriction enzyme site (**Figure S4B**). Among the total of 2,767 hygromycin-resistant callus lines analyzed, no CAPS positive callus lines were obtained.

**Figure 2.**
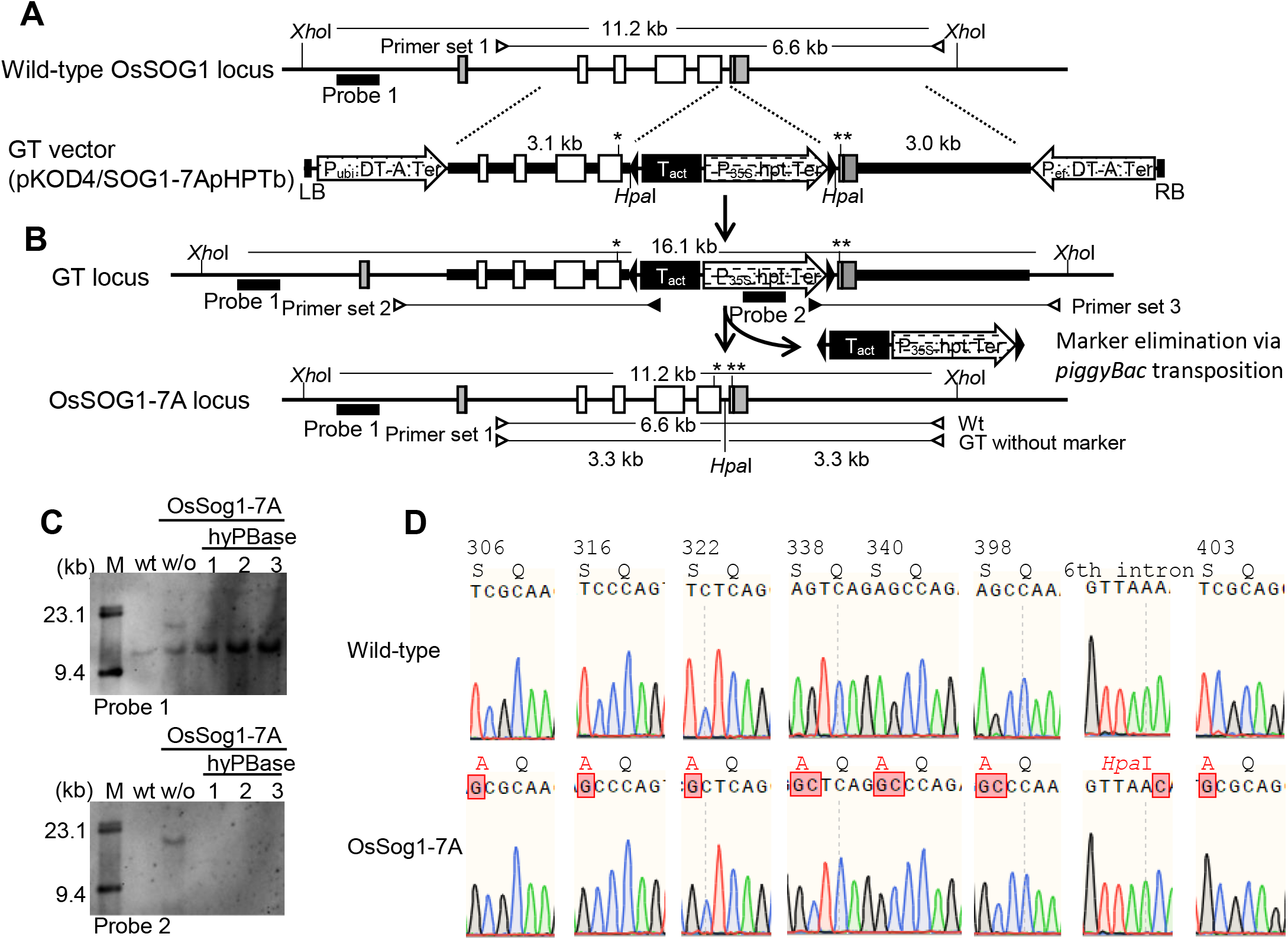
Generation of OsSOG1-7A mutants via GT and subsequent marker elimination using *piggyBac* transposon. A, Strategy for the introduction of point mutations into the *OsSOG1* locus via GT with positive-negative selection. The top line indicates the genomic structure of the wild-type *OsSOG1* locus. Gray and white boxes show untranslated regions and exons, respectively. The bottom line shows a schematic of the GT vector carrying the *DT-A* and *hpt* expression cassettes as a negative and positive selection markers, respectively. A rice actin terminator and *hpt* expression cassette (P_35S_:hpt:ter) were inserted within *piggyBac* transposon inverted repeats (black arrowheads) and introduced into the artificially synthesized *HpaI* site at the 5th intron of the sequence homologous to the *OsSOG1* locus. The sequence homologous to the *OsSOG1* locus on the GT vector carried a substitution of seven amino acids [S306A/S316A/S322A/S338A/S340A/S398A mutations at the 5th exon (*) and S403A mutation at the 6th exon (**)]. LB, left border; RB, right border. B, Strategy for marker excision from the GT locus via *piggyBac* transposition. The top line represents a schematic of the GT locus resulting from HR between the GT vector and wild-type locus. The bottom line shows the genomic structure of the OsSOG1-7A mutant locus containing the substitutions of seven amino acids and silent mutation (*Hpa*I site) resulting from precise marker elimination by *piggyBac* transposition. Primer set 1 was used for CAPS analysis to check the marker excision from the GT locus. C, Southern blot analysis with wild-type, OsSOG1-7A T_0_ plants transformed with (hyPBase-1, −2, and −3) or without (w/o) *piggyBac* transposase (hyPBase) expression cassette. Genomic DNA was digested with *XhoI.* Upper and lower panels were detected with probes 1 and 2, respectively, shown in A and B. D, Sequencing chromatograms of the substitution mutations of seven amino acids and the marker excision (*Hpa*I) site in wild-type (upper) and OsSOG1-7A T_2_ homozygous mutant (lower) plants.

The *OsSOG1* and *OsSGL* double mutant rice (termed *sog1sgl* mutant) were generated by CRISPR/Cas9-mediated mutagenesis to target the fourth exon of the *OsSOG1* and *OsSGL* genes using a gene-specific guide RNA (gRNA) (**Figure 3 A–C**). We selected two independent lines, *sog1sgl* #1, which contained a homozygous 25 bp deletion and homozygous 1 bp insertion in the *OsSOG1* and *OsSGL* genes, respectively, and #2, which carried a homozygous 1 bp deletion and homozygous 1 bp insertion in the *OsSOG1* and *OsSGL* genes, respectively, in the T_1_ generation (**Figures 3D and 3E**). These frameshift mutations in *sog1sgl* #1 and #2 mutants led to an early stop codon prior to the conserved SQ motifs in the C-terminal region of *OsSOG1* and *OsSGL* genes (**Figures 3D,E and S1**). In *sog1sgl* #1, CRISPR/Cas9-mediated mutations at the *OsSOG1* gene did not affect transcript levels, while full-length transcripts of *OsSGL1* were barely detectable by RT-PCR, probably due to nonsense-mediated mRNA decay (**Figure S3**). In addition, we confirmed by PCR analysis that these mutants lacked T-DNA. Their T_2_ or T_3_ progenies were subjected to further analysis.

**Figure 3.**
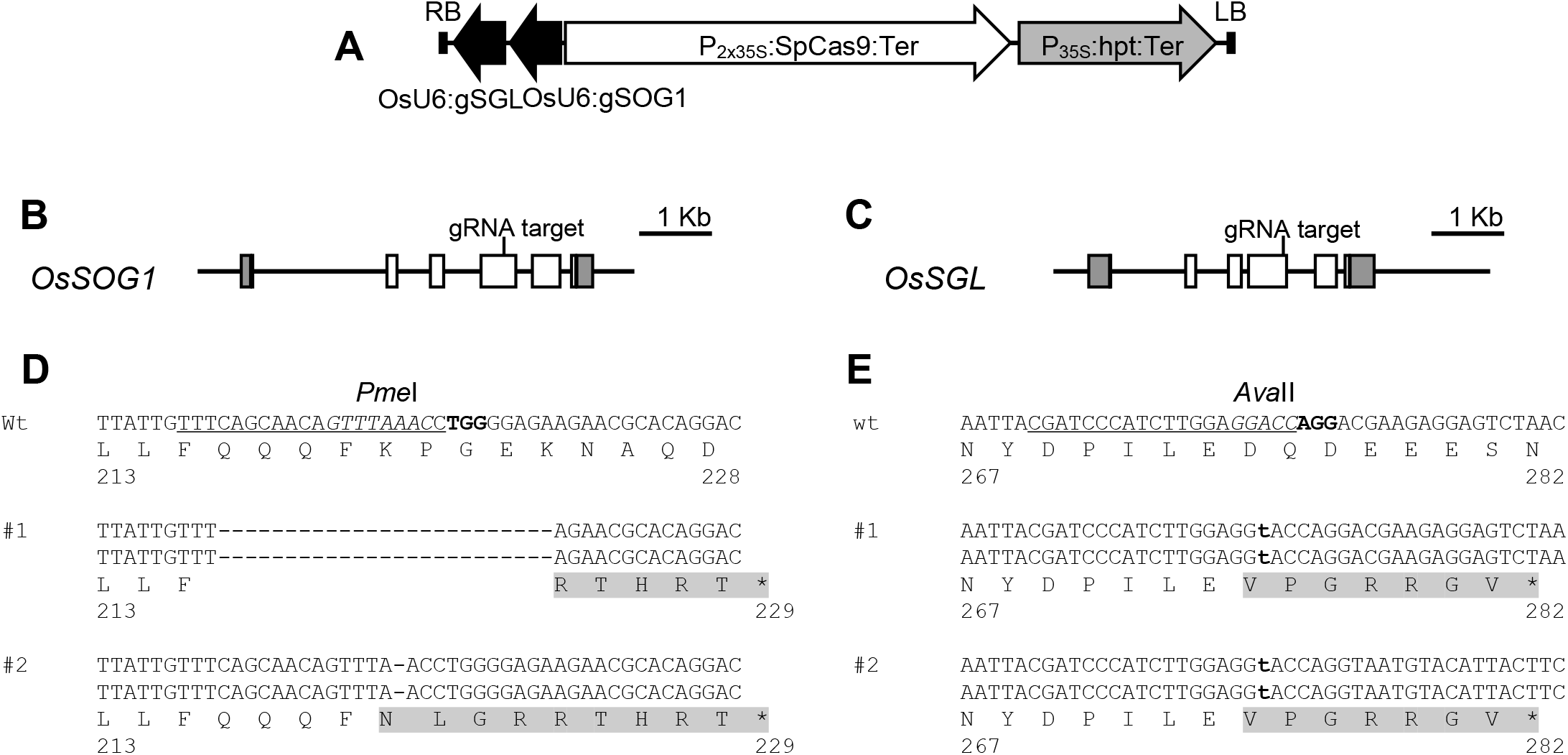
CRISPR/Cas9 target sites at OsSOG1 and OsSGL genes in *sog1sgl* double mutants. A, Construction of a CRISPR/Cas9 expression vector carrying the expression cassettes of the single gRNA targeting the *OsSOG1* (U6:gSOG1) and *OsSGL* (U6:gSGL) genes. B and C, Target sites of CRISPR/Cas9-mediated target mutagenesis in the *OsSOG1* (B) and *OsSGL* (C) genes. Gray and white boxes show untranslated region and exons, respectively. D and E, Mutation patterns induced via CRISPR/Cas9 system at *OsSOG1* (D) and *OsSGL* (E) gene in *sog1sgl* double mutants #1 and #2, respectively. *Pme*I (D) and *Ava*II (E) sites are shown in italics. Gray highlights indicate the deduced amino acid sequences in these mutants.

### OsSOG1 mutant and *sog1sgl* double mutant plants, but not OsSGL mutants, are highly sensitive to DNA damage agents

To elucidate the role of OsSOG1 and OsSGL in DDR and DNA repair in rice, we assessed the sensitivity of OsSOG1 and OsSGL mutants to the DNA damage agent bleomycin. Seeds of Psog1-GUS, Psgl-GUS, OsSOG1-7A mutants, and *sog1sgl* double mutants were sown on medium containing various concentrations of bleomycin and cultured for 1 week. Compared with that of wild-type plants, root growth of Psog1-GUS mutants was significantly inhibited by 10 μM bleomycin treatment (**Figure 4A and 4B**). However, the sensitivity of Psgl-GUS mutant plants was comparable with that of wild-type plants even under treatment with high concentration bleomycin. Although there was no statistically significant difference between the DNA damage susceptibility of OsSOG1-7A mutants and wild-type plants under treatment with 10 μM bleomycin, OsSOG1-7A mutants appeared to be more sensitive to DNA damage than wild-type or Psgl-GUS mutant plants. Inhibition of root growth of *sog1sgl* double mutants was observed at low concentrations of bleomycin, indicating that *sog1sgl* double mutants revealed the highest sensitivity among mutant plants used in this study. These results suggest that OsSOG1, but not OsSGL, works in DNA repair and/or DDR in rice, and its functions are partially regulated by ATM- and/or ATR-mediated phosphorylation of SQ motifs.

**Figure 4.**
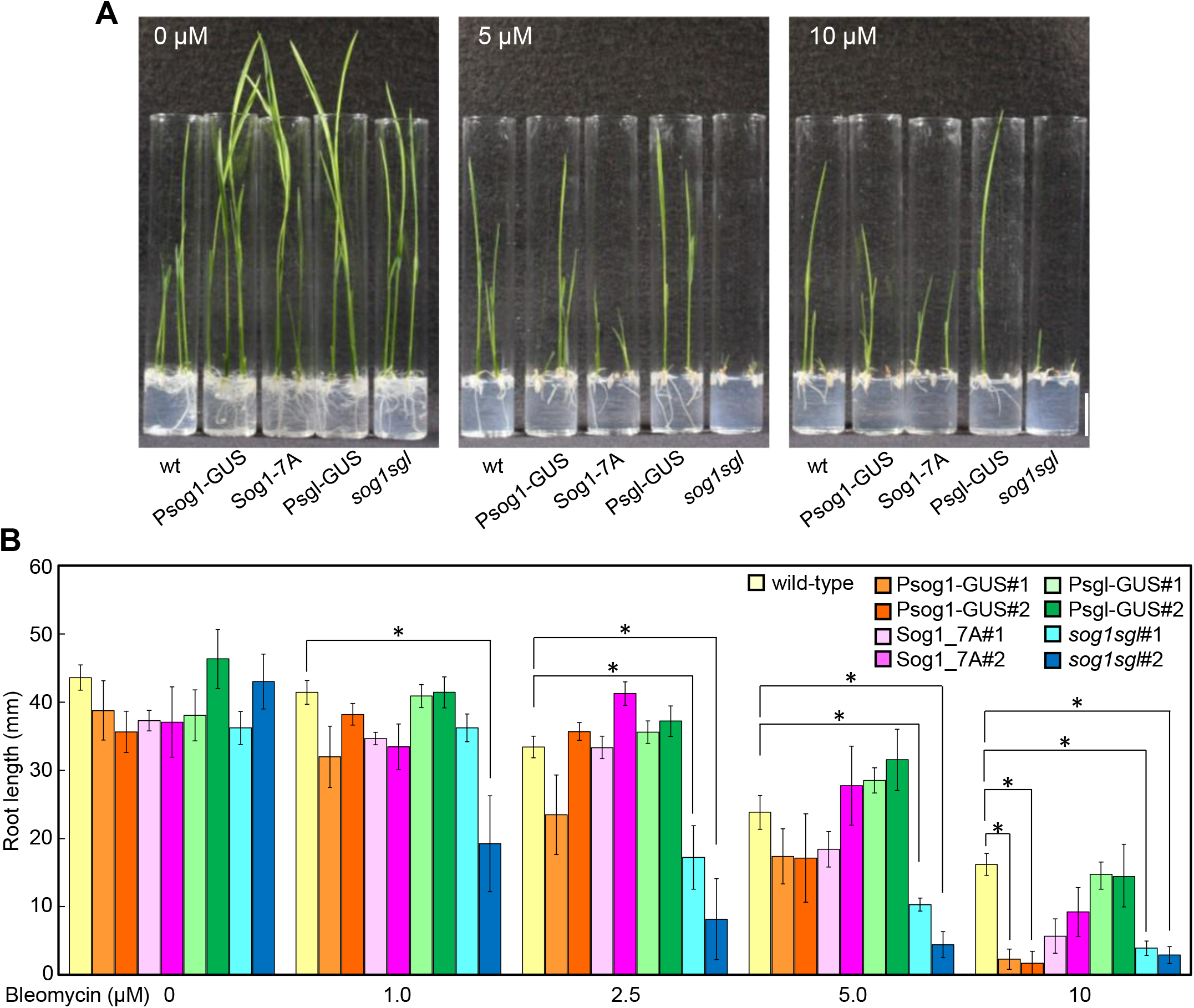
The effect of mutation in *OsSOG1* and/or *OsSGL* on sensitivity to the DNA-damaging agent Bleomycin. A, Representative morphology of 1-week-old wild-type, Psog1-GUS #1, Sog1-7A #1, and Psgl-GUS #1 sog1sgl #1 mutant rice plants grown with or without the indicated concentration of bleomycin. Bar=2cm. B, Quantification of the root length of plants grown with or without various concentrations of bleomycin. Error bars represent SE. T-test *, P<0.01.

### Analysis of DNA damage-response and tissue-specific expression of OsSOG1 and OsSGL in rice

Next, we analyzed transcript levels of *OsSOG1* and *OsSGL* genes in wild-type rice plants treated with bleomycin using quantitative RT-PCR (qRT-PCR). Transcript levels of OsSOG1 were barely affected by bleomycin treatment in either roots or shoots (**Figure S5A**) whereas transcript levels of *OsSGL* and *OsRad51A2—*well-known as DNA damage-response genes—increased significantly in response to bleomycin treatment, especially in roots (**Figure S5A**). Consistent with these data, GUS gene expression was detected in root tips of Psgl-GUS mutant plants upon treatment with bleomycin (**Figure S5B,C**). However, Psog1-GUS mutant plants displayed no detectable GUS expression either with or without bleomycin treatment. Based on these results, microarray analysis was performed with roots to evaluate the effect of mutation of OsSOG1 on the expression of genes responsive to DNA damage.

### Transcriptome analysis in OsSOG1 and OsSGL mutants under treatment with a DNA damage agent

To identify target genes regulated by OsSOG1 and/or OsSGL under DNA damage conditions, 1-week-old seedlings of wild-type and Psog1-GUS, Psgl-GUS, OsSOG1-7A mutants, and the *sog1sgl* double mutant were treated with or without 10 μM bleomycin for 8 h and harvested for microarray analysis. Total RNA extracted from roots (about 2 cm area from the root tip) was subjected to microarray analysis. Microarray analysis using wild-type with and without bleomycin treatment showed that the transcript levels of 1,068 probes were induced (**Table S1**) and 608 probes were reduced by more than two-fold (*P* < 0.01) in response to bleomycin treatment (**Table S2**). In Psog1-GUS, OsSOG1-7A and *sog1sgl* mutants, the number of probes upregulated in response to DNA damage was only about half of that in wild-type (**Table 1**). On the other hand, mutation of *OsSGL* was thought not likely to affect gene expression profiles in response to DNA damage.

To define the downstream genes of OsSOG1-mediated DDR in rice, we identified the probes upregulated or downregulated more than two-fold in wild-type treated with bleomycin compared with non-treated controls, as well as transcriptional changes reduced to less than half of wild-type in *sog1* mutant plants even under DNA damage conditions. Within the 1,068 bleomycin-induced probes in wild-type, induction of 758 and 738 probes was canceled in both of two independent lines of Psog1-GUS #1 and #2, or OsSOG1-7A #1 and #2, respectively (**Table S1**). Among these, 608 probes were consistently not induced in four lines of Psog1-GUS#1, #2, OsSOG1-7A#1, and #2 compared with wild-type (**Figure 5A**). The average fold changes of 60% (362/608) probes were higher in OsSOG1-7A#1, #2 than in Psog1-GUS#1, #2, suggesting the persistence of transcriptional activity of the OsSOG1-7A mutant. The induction of 908 probes in response to bleomycin treatment was impaired in the *sog1sgl* mutant (**Figure 5A**). Loss of the *OsSGL* gene led to a decrease in the 278 probes responsive to DNA damage in Psgl-GUS mutants #1 and #2 (**Table S1 and Figure 5B)**. Among 608 downregulated probes in response to bleomycin in wild-type, no reduction in the levels of 237, 417, and 192 probes was found in either of two independent lines of Psog1-GUS, OsSOG1-7A, or Psgl-GUS, respectively (**Table S2, Figure 5C, and D)**. The downregulated 501 probes in wild-type treatment with bleomycin were not reduced in *sog1sgl* mutant following treatment with bleomycin.

**Figure 5.**
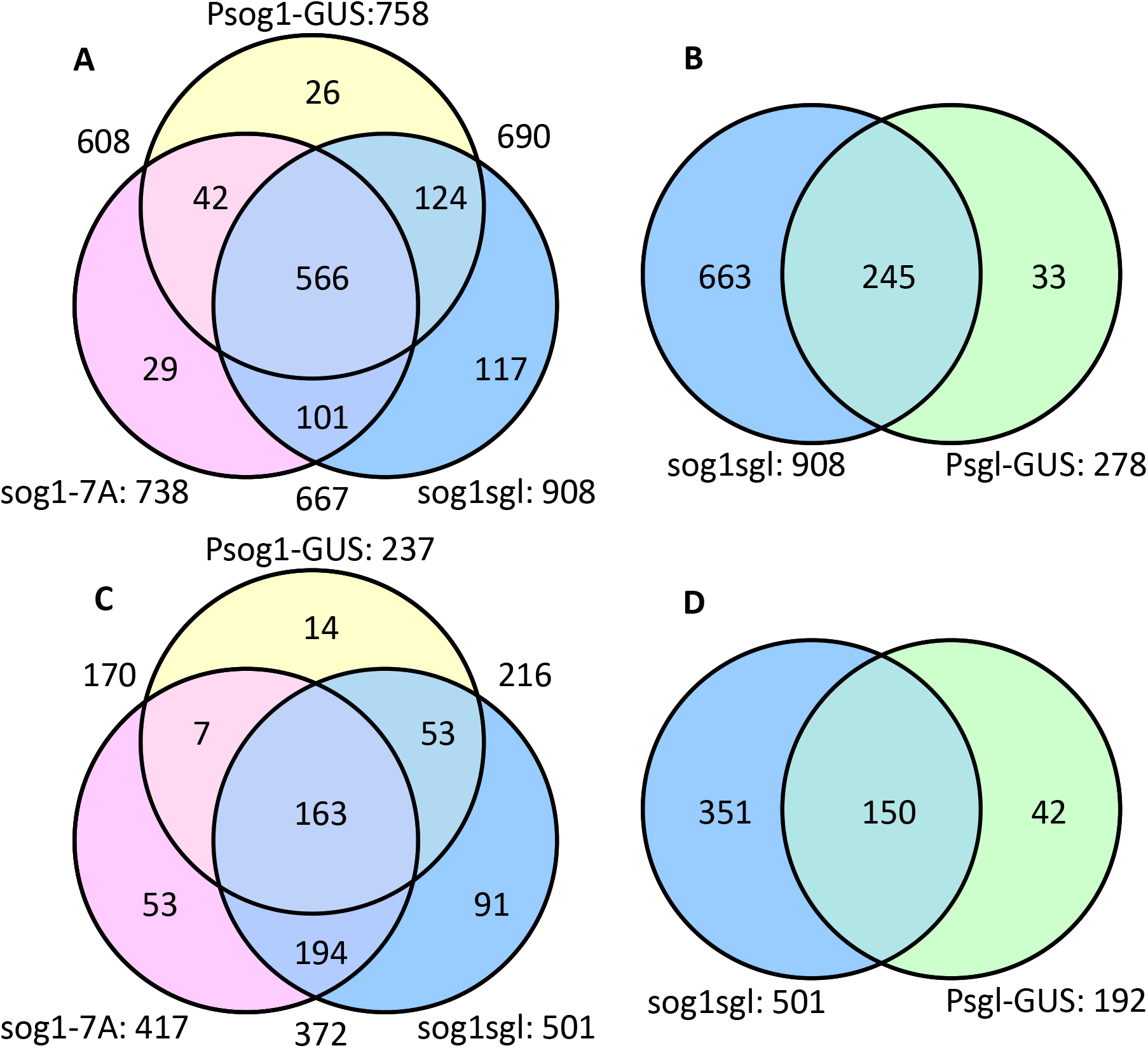
Identification of *OsSOG1* and *OsSGL* target genes by microarray analysis. A and B, Venn diagrams showing the overlap of microarray probes that were repressed in Psog1-GUSs/OsSOG1-7As/*sog1sgl*s (A), and in *sog1sgl*s/Psgl-GUSs (B), respectively, among the bleomycin-induced 1,068 probes in wild-type. C and D, Venn diagrams representing the overlap of microarray probes that were not reduced in Psog1-GUSs/OsSOG1-7As/*sog1sgl*s (C), and in *sog1sgl*s/Psgl-GUSs (D), respectively, among 608 probes downregulated in response to bleomycin in wild-type rice.

The bleomycin-induced 566 and bleomycin-reduced 163 probes in wild-type were consistently not reduced in Psog1-GUS#1, #2, OsSOG1-7A#1, #2, and *sog1sgl* mutant plants (**Figure 5A and 5C**). However, it is thought that OsSOG1-7A mutants led to partial disruption of OsSOG1 function based on the persistence of transcriptional activity and DNA damage susceptibility of OsSOG1-7A mutants; thus, we defined 690 bleomycin-induced and 216 bleomycin-reduced probes in which expression levels were not changed in Psog1-GUS and *sog1sgl* mutant plants as candidate target genes of OsSOG1-mediated DDR (**Table S1**). Gene ontology (GO) enrichment analysis with ShinyGO (v0.66, http://bioinformatics.sdstate.edu/go/) revealed that “Glutathione metabolic process”, “Meiotic cell cycle process”, “Double-strand break repair”, “Homologous recombination”, and “Jasmonic acid metabolic process” etc., were enriched in upregulated OsSOG1-target candidate genes in response to bleomycin treatment (**Table S3**). Functional categories regarding chromatin organization were concentrated in the bleomycin-reduced OsSOG1-target candidate gene (**Table S4**).

The OsSOG1-target candidate contained some transcription factors including OsSGL, indicating that these genes comprised genes regulated directly and indirectly by OsSOG1. It has been reported previously that *Arabidopsis* SOG1 binds to the CTT (N)_7_ AAG motif on the promoter region of AtSOG1 target genes (Ogita et al., 2018). We validated whether CTT (N)_7_ AAG motifs were located within 1 kb upstream of the start site of the protein-coding region (CDS) of upregulated and downregulated transcripts in wild-type in response to treatment with bleomycin. As a result of the alignment between the microarray probe sequences and the reference genome sequence (the Nipponbare IRGSP-1.0 reference genome), the 785 upregulated- and 477 downregulated-transcripts among 1,068 upregulated- and 608 downregulated-probes in wild-type treated with bleomycin, respectively, were used to search for the AtSOG1 binding motif in the promoter region. AtSOG1 binding motifs with the sequence CTT (N)_7_ AAG were found in the 1-kb region upstream of the start site of the CDS of 337 bleomycin-induced (337/785, 42.9%, **Table S5**) and 87 bleomycin-reduced transcripts (87/477, 18.2%, **Table S6**). Among them, 74.8% (252/337, **Table S5**) and 28.7% (25/87, **Table S6**) bleomycin-induced and reduced transcripts were contained in the OsSOG1-target candidate, indicating that these are regulated directly by OsSOG1.

### OsSOG1 regulated the expression of DNA repair-related genes directly in response to DNA damage

Microarray analysis showed that the bleomycin-induced OsSOG1-target candidates included DNA repair-related genes, such as OsRad51A2, OsMSH4, OsMSH5, OsDMC1A, DMC1B, Tyrosyl-DNA phosphodiesterase family protein, Regulator of telomere elongation helicase, Rad21/Rec8 family protein, and Kleisin Nse4 domain-containing protein, etc. To confirm these data, we analyzed the expression levels of representative DNA repair-related genes by qRT-PCR. The expression levels of all genes induced by treatment with bleomycin (Rad51A2, PARP2A, DMC1A, DMC1B, MSH4, MSH5, Rad21-2, Rad21-4, and BRCA1) were suppressed completely in Psog1-GUS and *sog1sgl* mutants (**Figure 6**). In Sog1-7A mutants, the expression levels of these DNA repair-related genes were also depressed, even upon treatment with bleomycin, although often not completely. These results provided support for the data that SOG1-7A mutation has a partial transcriptional activity revealed by the microarray analysis whereas mutation of OsSGL did not affect induction of these DNA repair-related genes.

**Figure 6.**
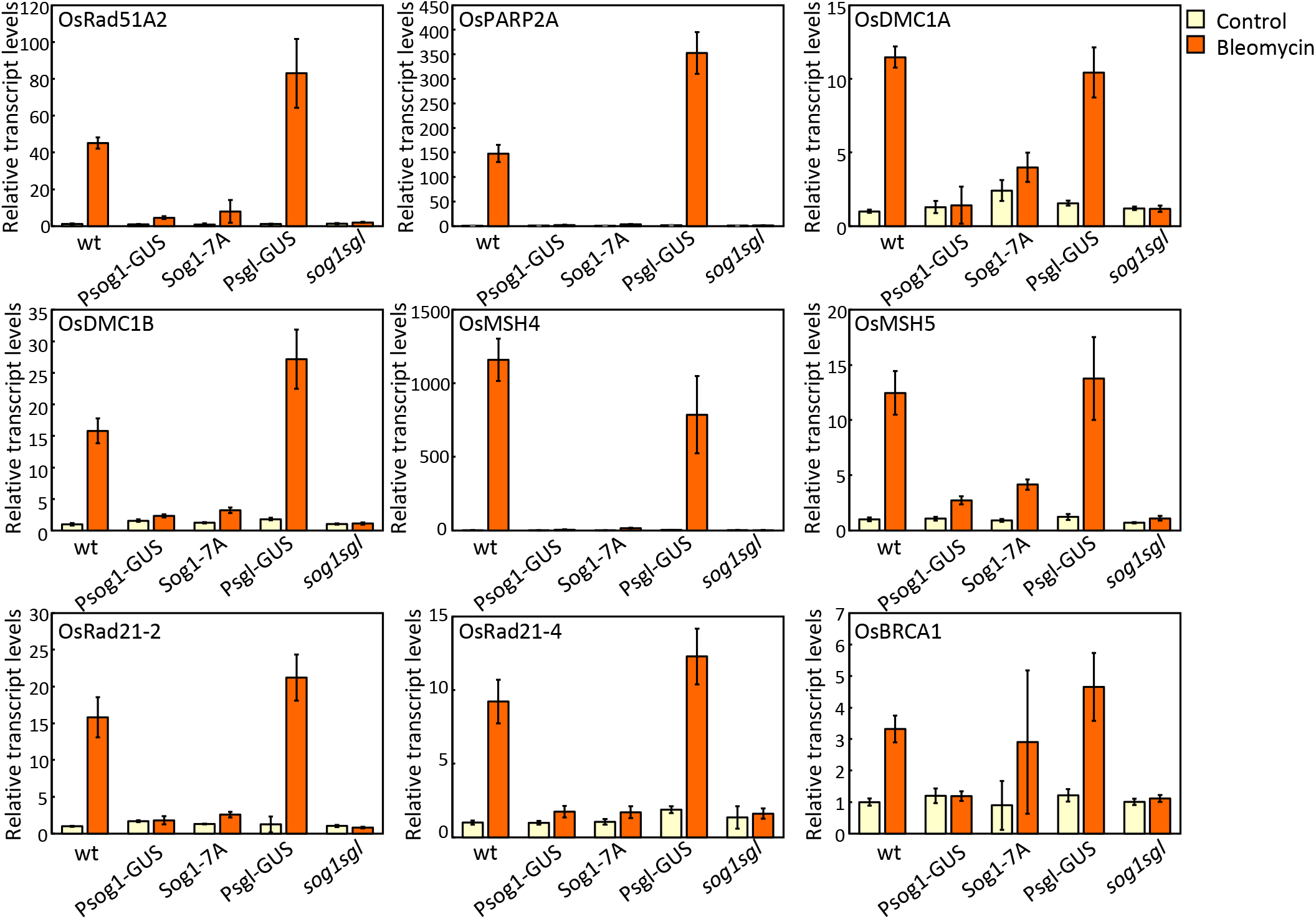
Transcript levels of DNA repair-related genes in response to DNA damage in wild-type, Psog1-GUS, Sog1-7A, Psgl-GUS, and *sog1sgl* mutant plants. Plants were treated with 10 μM bleomycin for 8 h, and roots were harvested and subjected to RNA extraction and qRT-PCR analysis.

AtSOG1 binding motifs [CTT (N)_7_ AAG] were found on the promoter of almost all DNA repair-related genes that were upregulated in response to DNA damage treatment (**Table S5**). To assess whether the AtSOG1 binding motif mediated the upregulation of these DNA repair-related genes in response to DNA damage, we introduced mutations in the AtSOG1 binding motif on the promoter of OsRad51A2 and OsDMC1A via CRISPR/Cas9-based mutagenesis (**Figure 7A,B**). Wild-type rice calli were transformed with expression vectors of CRISPR/Cas9 with a gRNA expression cassette targeting the CTT(N)_7_-AAG motif on the promoter of these genes via *Agrobacterium-mediated* transformation. Transgenic calli were propagated clonally and subjected to DNA heteroduplex mobility assay (HMA) to detect CRISPR/Cas9-mediated targeted mutagenesis at the target site. Transgenic plants were regenerated from callus lines harboring mutations at high and low frequency at the target site, and plants with and without CRISPR/Cas9-mediated targeted mutations were then isolated using sequencing analysis. Plants with or without mutations at the CTT(N)_7_AAG motif on the promoter of OsRad51A2 or OsDMC1A were treated with bleomycin and collected for analysis of the bleomycin-inducible expression of these genes. The expression levels of OsRad51A2 or OsDMC1A were significantly induced in response to bleomycin treatment in control plants [without mutations by CRISPR/Cas9 at CTT(N)_7_AAG motif]; however, the induction of bleomycin-responsible expression was impaired in mutant plants. These results suggest that the DNA damage-inducible expression of DNA repair-related genes, OsRad51A2 and OsDMC1A, was regulated via the CTT(N) 7AAG motif in rice as in Arabidopsis.

**Figure 7.**
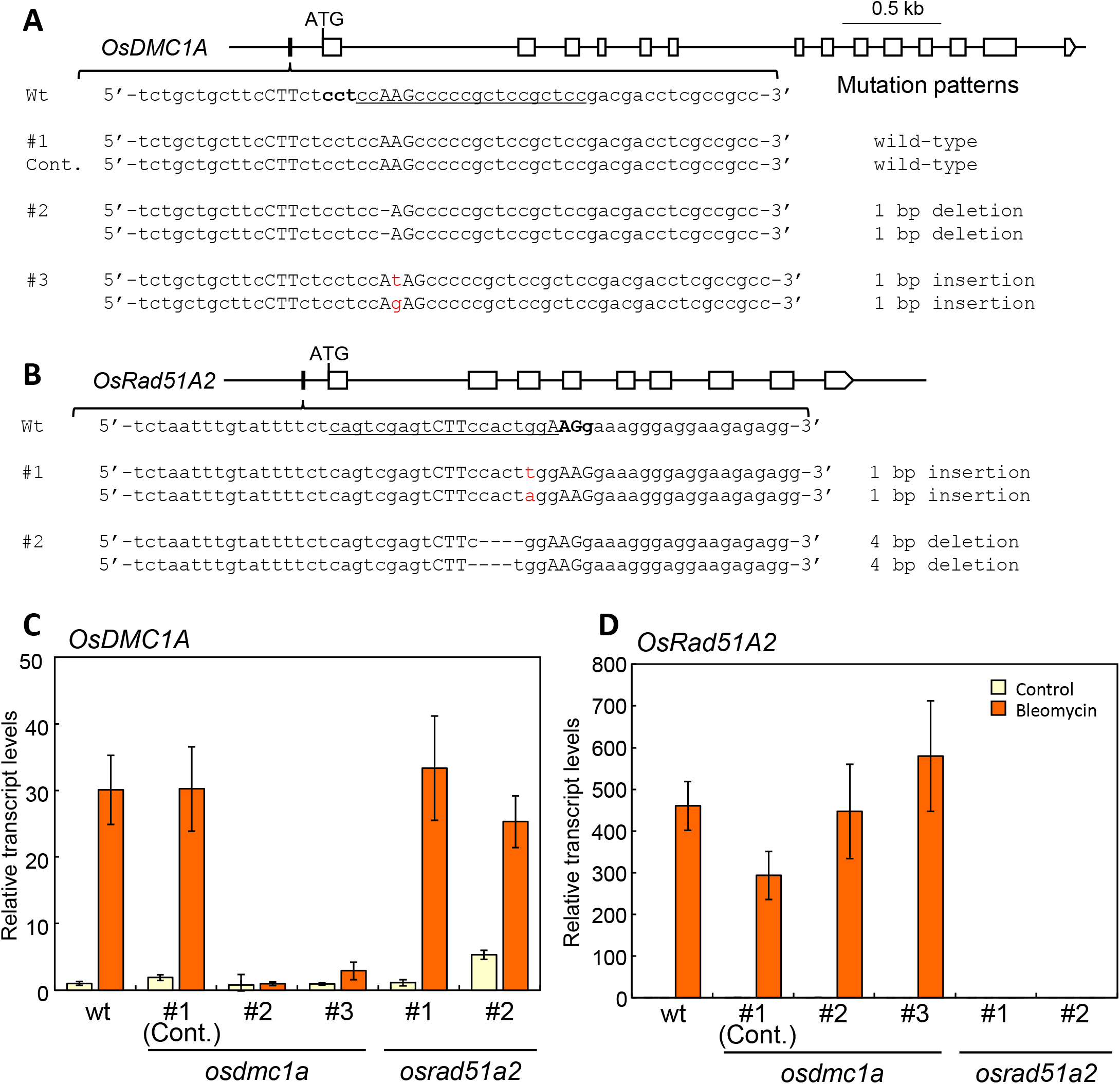
CTT(N)_7_AAG motif-mediated regulation of the *OsDMC1A* and *OsRad51A2* genes by *OsSOG1*. A and B, Mutation patterns induced via the CRISPR/Cas9 system at CTT(N)_7_AAG motifs (uppercase letters) on the promoter region of *OsDMC1A* (A) and *OsRad51A2* (B) genes in *osdmc1a* and *osrad51a2* mutant plants. The top lines indicate the genomic structures of *OsDMC1A* (A) and *OsRad51A2* (B) genes, respectively. Solid and open boxes represent CTT(N)_7_AAG motifs and open reading frames, respectively. The underlined sequence indicates the 20 bp target sequence for the sgRNA, and bold letters indicate the PAM sequence. The dashes and red letters show deletion and insertion mutations. The osdmc1a #1 was used as control that did not have mutations at CTT(N)_7_AAG motifs even with the CRISPR/Cas9 vector. C and D, Transcript levels of *OsDMC1A* and *OsRad51A2* in wild-type, *osdmc1a*, and *osrad51a2* mutant plants treated with or without 10 μM bleomycin.

### Some cell cycle-related genes are regulated by OsSOG1

A previous study demonstrated that AtSOG1 causes G2 arrest in the meristem through regulation of CDK inhibitor expression under DNA damage conditions (Adachi et al., 2011). Therefore, we analyzed the expression levels of cell cycle-related genes and revealed significant expression changes in response to DNA damage in microarray using qRT-PCR. The expression level of EL2—a cyclin-dependent kinase inhibitor—was induced significantly in response to bleomycin treatment in wild-type; however, its induction was cancelled in Psog1-GUS mutants (**Figure S6**). Bleomycin treatment led to downregulation of CDKB2;1 and CYCB2;1 expression in wild-type but not in Psog1-GUS mutants, whereas there was no significant difference in the expression of KNOLLE, CYCA1;1, CYCA1;2, CYCA3;2, CYCB1;1, CYCB1;2 and CYCB2;2 between wild-type and Psog1-GUS mutants (**Figure S6**).

### OsSOG1 plays an important role not only in the DNA damage but also the oxidative stress response

In Arabidopsis, AtSOG1 is involved not only in DDR but also in the oxidative stress response under abiotic stress conditions such as exposure to cadmium, aluminium, and UV-B light. Treatment with bleomycin induced the expression of ROS homeostasis-related genes (e.g., glutathione-S-transferase and dehydroascorbate reductase) in rice, and its bleomycin-inducible expression was suppressed in the OsSOG1 mutant. To test the hypothesis that OsSOG1 can contribute to oxidative stress tolerance by regulation of expression of ROS homeostasis-related genes, we evaluated the sensitivity of OsSOG1 mutant plants to the herbicide methyl viologen (MV), which produces ROS in the chloroplast under illumination. All sog1 mutant plants (Psog1-GUS, SOG1-7A, and *sog1sgl* mutant) showed increased susceptibility to MV compared with wild-type plants (**Figure S7**).

## Discussion

A database search identified OsSOG1 and OsSGL as potential rice orthologs of the Arabidopsis DDR protein AtSOG1. The functional SQ motif of AtSOG1 was conserved completely or partially [Aligned score: 47.1292 (AtSOG1 vs OsSOG1) and 39.1852 (AtSOG1 vs OsSGL)] and DNA damage susceptibility tests and transcriptome analyses using OsSOG1 and OsSGL mutant rice demonstrated that deficiency of OsSOG1, but not of OsSGL, led to significantly increased sensitivity to DNA damage and inhibition of the induction of gene expression in response to DNA damage. Because induction of OsSGL expression in response to DNA damage was repressed in OsSOG1 mutants, and a SOG1 binding motif was identified in the promoter region of OsSGL, OsSGL was thought to be regulated directly by OsSOG1. However, *sog1sgl* double mutant plants showed a more severe phenotype to DNA damage and a stricter suppression of DDR gene expression than that of the *OsSOG1* single mutant Psog1-GUS. Thus, OsSGL might have a role not only as a downstream target of OsSOG1 but also as a member of another signal transduction pathway.

Phosphorylation of AtSOG1 via ATM is reported to be required for the DDR in Arabidopsis (Yoshiyama et al., 2013, 2017). Expression of AtSOG1 with serine-to-alanine substitutions (that is, non-phosphorylatable mutations) at all five SQ motifs in the *Arabidopsis sog1* mutant does not complement the sog1 mutant phenotype at all (Yoshiyama et al., 2017), whereas a recent study demonstrated that casein kinase 2 (CK2) also regulated AtSOG1 activity through phosphorylation in response to DNA damage conditions (Wei et al., 2021). In the present study, we used GT to generate OsSOG1-7A mutants, which carried substitutions of serine to alanine at all seven SQ motifs of OsSOG1, and observed that these mutants were less sensitive to bleomycin than Psog1-GUS and *sog1sgl* mutant plants (**Figure 4**), indicating that the substitution SQ motif mutations of OsSOG1 inhibited OsSOG1 activity, but not completely. The OsSOG1 protein has nine potential CK2 phosphorylation sites [(S/T)XX(E/D)], raising the possibility that OsSOG1 is also regulated via CK-mediated signaling in a DDR pathway. Although we attempted to introduce phosphor-mimetic mutations (the substitution of serine with aspartic acid) at SQ motifs of OsSOG1 by CRISPR/Cas9-mediated GT (developed in rice by Nishizawa-Yokoi et al., 2020), mutant plants carrying desired mutations in *OsSOG* (OsSOG1-7D mutants) were not obtained using this method (**Figure S4**). Unlike GT with positive-negative selection, DSBs were induced at the *OsSOG1* locus via CRISPR/Cas9 using CRISPR/Cas9-mediated GT; therefore, one allele was disrupted by the introduction of insertion or deletion mutations at the CRISPR-target site even if GT had occurred on another allele. This could explain why we could not obtain phospho-mimetic mutants (OsSOG1-7D) rice. There is much room for examination regarding the mechanisms of OsSOG1 activation in response to DNA damage.

Microarray analysis and qRT-PCR revealed that DNA repair-related genes were regulated by OsSOG1 in response to DNA damage in a manner similar to that of AtSOG1 reported previously (Yoshiyama et al., 2009; Ogita et al., 2018). Rad51, BRCA1, and PARP2—representative DNA damage-response genes—were identified as SOG1 target genes in both Arabidopsis and rice. However, OsMSH4, OsMSH5, OsDMC1A, OsDMC1B, OsRad21-2, and OsRad21-4 genes were isolated as OsSOG1-specific target genes (**Table S1 and Figure 6**). MSH4 and MSH5 encode a meiosis-specific member of the MutS-homologous family of genes, and these proteins have been revealed to interact with each other and to promote the formation of crossovers between homologous chromosomes in meiosis both in Arabidopsis and rice (Higgins et al., 2008; Luo et al., 2013; Zhang et al., 2014). Similar to Rad51, DMC1 is a eukaryotic RecA homolog. It has been reported that both Rad51 and DMC1 bind to the single-stranded DNA processed from DSBs and promote single-strand invasion to search for homologous sequences during HR in many species, including Arabidopsis and rice (Bishop et al., 1992; Shinohara et al., 1997; Kobayashi et al., 2018). In various eukaryotes, it is thought that Rad51 is involved in both mitotic and meiotic HR, whereas DMC1 catalyzes specifically meiotic HR (Shinohara et al., 1997; Kurzbauer et al., 2012; Pradillo et al., 2012; Da Ines et al., 2013). Wang et al. (2016) reported that *Osdmc1a Osdmc1b* double mutant rice plants were completely sterile because of meiotic defects (Wang et al., 2016). Rad21, and its meiosis-specific variant Rec8, have been reported to play a significant role in sister chromatid cohesion and segregation for mitosis and meiosis, respectively, in yeast (Watanabe and Nurse, 1999). Among four Rad21/Rec8 homologs encoded in the rice genome (Zhang et al., 2006), at least OsRad21–4 functions in homologous pairing of chromosome and telomere bouquet formation during the meiotic processes (Shao et al., 2011). Considering that these factors were upregulated via OsSOG1 in somatic cells under DNA damage conditions, it seems likely that they are involved in DNA repair, especially HR, during mitosis.

It has been reported that DNA damage inhibits plant cell division and induces endoreplication, in which cells increase their genomic DNA content without dividing, in *Arabidopsis* (Adachi et al., 2011). Arabidopsis SOG1 has been reported to be involved in both inhibition of the cell cycle and induction of the endocycle during DNA damage (Yoshiyama et al., 2009; Adachi et al., 2011; Yoshiyama, 2016; Ogita et al., 2018). In contrast to *Arabidopsis*, endocycle has not been found in rice other than during endosperm development, even under stress conditions (Endo et al., 2012). Cell cycle arrest occurs via transcriptional activation of SIAMESE/SIAMESE-RELATED 5 and 7 (SMR5 and SMR7), cyclin-dependent kinase inhibitors, with a subsequent decrease in the expression of cyclin A (CYCA) and B (CYCB) genes, which are essential for CDK activation at the G2/M transition of the mitotic cell cycle, and KNOLLE, which is required for cytokinesis, depends on SOG1 in Arabidopsis (Adachi et al., 2011; Yi et al., 2014; Chen et al., 2017, 2019). Thus, root elongation of Arabidopsis *sog1* mutants was not inhibited on medium containing DNA damage agents, resulting in a more tolerant phenotype of Arabidopsis *sog1* mutants to DNA damage compared with wild-type (Yoshiyama et al., 2009, 2014; Sjogren et al., 2015). However, in the present study, root growth of the rice *sog1* mutant was suppressed on medium containing bleomycin at any concentration in comparison to wild-type (**Figure 4**). Microarray analysis and qRT-PCR revealed that expression of EL2, a cyclin-dependent kinase inhibitor, increased significantly in response to DNA damage in wild-type, but this induction was lost in rice *sog1* mutants (**Figure S6**). In addition, downregulation of OsCDKB2;1 and OsCYCB2;1 in response to DNA damage likely depended on OsSOG1; however, the regulation of expression of many other cell cycle-related genes (e.g. OsKNOLLE, OsCYCB1;1 and OsCYCB1;2) was independent of OsSOG1 (**Figure S6**). Surprisingly, no transcriptional change of rice cyclin-dependent kinase inhibitors, other than EL2, was found even after bleomycin treatment in wild-type.

These results suggest that the mechanisms by which cells prevent DNA damage in rice, especially regarding the induction of cell-cycle arrest and endocycle, might differ from those of *Arabidopsis*. Such differences could be attributed to the distribution of dividing cells within the body of the plant. In contrast to Arabidopsis, which has dividing cells throughout its whole structure, dividing cells exist only in meristematic tissue in rice. As mentioned above, OsSOG1-targeted DNA repair factors, e.g. OsDMC1s and OsMSH4/5, which were thought to be meiosis-specific recombination proteins in many organisms, might be involved in somatic HR during DNA damage in rice, raising the possibility that DNA repair, especially HR, is strengthened in somatic cells under DNA damage conditions. This phenomenon is assumed to be a rice-specific DDR. Further investigations are required to clarify the mechanisms of DDR in rice.

In Arabidopsis, genes involved in defense responses, to not only abiotic stress but also biotic stress, include SOG1 target genes. In addition, *sog1* mutants exhibit decreased resistance to infection with certain types of fungus in Arabidopsis (Ogita et al., 2018). Furthermore, the flavin-containing monooxygenase (FMO1) gene related to activation of systemic acquired resistance through salicylate acid signaling was reported to be regulated directly by Arabidopsis SOG1 (Chen and Umeda, 2015). Under DNA damage conditions, OsSOG1 was shown by microarray analysis to regulate the expression of a large number of detoxifying enzymes, including glutathione-S-transferases (GSTs) and cytochrome P450 (CYPs). The DNA damage-inducible expression of ROS homeostasis-related genes and the content of glutathione in shoots and roots under DNA damage conditions were also affected by OsSOG1 mutation. In addition, microarray analysis revealed that jasmonic acid (JA) metabolic process-related genes were enriched in the OsSOG1 target. Many previous studies have reported that JA plays a crucial role in rice immunity against bacterial and fungal infection (Yamada et al., 2012; Ranjan et al., 2015; Ghorbel et al., 2021). These results suggested that OsSOG1 induces multiple defense pathways in response to DNA damage and plays a key role in acquiring resistance against DNA damage, as in Arabidopsis.

Recently developed genome editing technologies have become an essential tool not only for basic plant science but also for plant breeding. Although these techniques were categorized as targeted mutagenesis and HR-mediated GT, these mechanisms introduce mutations via the DSB repair pathways of NHEJ, and HR, respectively. Therefore, extending our knowledge of the response to DNA damage and the regulation mechanisms of DNA repair pathways is important for the establishment of efficient plant genome editing technologies. Elucidation of the regulatory mechanisms of OsSOG1-mediated DDR and DNA repair will contribute to the development of efficient and widely applicable genome editing methods via regulation of OsSOG1 activity in the future.

## Methods

### Plant materials

*Oryza sativa* L. cv. Nipponbare was used in this study. Rice calli were grown on N6D medium at 33°C under continuous light, and plants were regenerated from calli on ReIII medium or germinated from seeds grown at 28°C under 16 h light:8 h dark conditions.

### Vector construction

The GT vectors, pKOD4/Psog1-GUS_HPT (**Figure 1A**) and pKOD4/Psgl-GUS_HPT (**Figure S2A**) used to generate Psog1-GUS and Psgl-GUS mutant rice using positive-negative selection were constructed as follows. The hygromycin phosphotransferase (hpt) expression cassette containing the CaMV 35S terminator (T35S)::nopaline synthase terminator (Tnos)::CaMV35S promoter (P35S)::hpt gene::rice heat shock protein 17.3 terminator (Thsp17.3) was cloned into *AscI/PacI* site between attL1 and attL2 sequence in pENTR(L1-L2) (Thermo Fisher Scientific), yielding pE(L1-L2)hpt. The promoter regions (3 kb upstream from ATG) of the OsSOG1 and OsSGL genes and the GUS gene coding region were amplified by PCR from rice genomic DNA or pENTR-gus plasmid (Thermo Fisher Scientific), respectively, using the primer sets listed in **Table S7**. PCR fragments representing either the OsSOG1 promoter+GUS gene or OsSGL promoter+GUS gene were inserted into the *Asc*I/*Avr*II site using an In-Fusion HD cloning kit (Takara) between attL1 and T35S in pE(L1-L2)hpt, yielding pE(L1-L2)Psog1-GUS_hpt and pE(L1-L2)Psgl-GUS_hpt, respectively. The 3-kb DNA fragments with *OsSOG1* or *OsSGL* coding region were amplified by PCR from rice genomic DNA and cloned into the *Pac*I site between Thsp17.3 and attL2 in pE(L1-L2)Psog1-GUS_hpt and pE(L1-L2)Psgl-GUS_hpt, respectively. Finally, DNA fragments of the OsSOG1 promoter::GUS::T35S::Tnos::P35S::hpt::Thsp17.3::OsSOG1 coding region and OsSGL promoter::GUS::T35S::Tnos::P35S::hpt::Thsp17.3::OsSGL coding region were cloned into pKOD4 (Nishizawa-Yokoi et al., 2015) with two *DT-A* gene expression cassettes (maize polyubiquitin 1 promoter + DT-A + rice hsp 16.9a terminator and rice elongation factor-1a promoter + DT-A + rice hsp 16.9b terminator) by a Gateway LR clonase II reaction (Thermo Fisher Scientific), yielding pKOD4/Psog1-GUS_HPT (**Figure 1A**) and pKOD4/Psgl-GUS_HPT (**Figure S2A**), respectively.

The GT vector pKOD4/SOG1_7ApHPTb used to produce OsSOG1–7A mutants carrying a serine to alanine substitution at all seven SQ motifs of the OsSOG1 gene by positive-negative selection was constructed as follows. A 0.6 kb artificially synthesized fragment from *Bsr*GI to *Bsr*GI (Thermo Fisher Scientific) of the *OsSOG1* gene containing the 5th exon (with S306A/S316A/S322A/S338A/S340A/S398A mutations), 5th intron (with silent mutation producing *Hpa*I site, gttaaa to gttaac), 6th exon (with S403A mutation), and *3’* UTR region was cloned into the *BamHI/XhoI* site of pBluescript KS– using an In-Fusion HD cloning kit (Takara), yielding pOsSOG1_7A. The *piggyBac* inverted-repeat transposable element harboring the meganuclease I-*Sce*I site (Nishizawa-Yokoi et al., 2014) was cloned into a *HpaI* site in the 5th intron of *OsSOG1* in pOsSOG1_7A, yielding pOsSOG1_7Apb. A 2.6 kb upstream and 2.9 kb downstream fragments from two *BsrGI* sites at the 4th intron and 3’ UTR region of the OsSOG1 gene were amplified by PCR from rice genomic DNA and inserted together with the OsSOG1_7Apb fragment digested with *Bsr*GI into the *AscI/PacI* site of pENTR(L1-L2) (Thermo Fisher Scientific) using an In-Fusion HD cloning kit (Takara), yielding pE(L1-L2)OsSOG1_7Apb. A 4.3 kb positive selection marker expression cassette containing the rice actin terminator, P35S, hpt and Thsp17.3 was digested with I-*Sce*I and integrated into the I-*Sce*I site within the *piggyBac* inverted repeat of pE(L1-L2)OsSOG1_7Apb, yielding pE(L1-L2)OsSOG1_7ApHPTb. The OsSOG1_7ApHPTb fragment was re-cloned into the GT binary vector pKOD4 using a Gateway LR clonase II reaction (Thermo Fisher Scientific), yielding pKOD4/ SOG1_7ApHPTb (**Figure 2A**).

To construct the CRISPR/Cas9 expression vector targeting *OsSOG1* and *OsSGL* genes, 20-nt annealed oligonucleotide pairs for the target sequences shown in **Table S7** were cloned into the *BbsI* site of the sgRNA expression vector described as pU6gRNA-oligo (Mikami et al., 2015). The sgRNA expression cassette targeting OsSGL (OsU6::sgOsSGL) was digested with *PvuII/AscI* and ligated into the *Eco*RV/*Asc*I sites in pU6::sgOsSOG1 with the sgRNA expression cassette targeting the OsSOG1 gene to connect these two sgRNA expression cassettes in tandem. The fragment with two sgRNA expression cassettes (OsU6::sgOsSOG1 and OsU6::sgOsSGL) was digested with *Asc*I/*Pac*I and introduced into pZH_OsU6sgRNA_MMCas9 (Mikami et al., 2015). The CRISPR/Cas9-mediated GT vector used to introduce phosphomimetic mutations (substitutions of serine to aspartic acid) at all seven SQ motifs of *OsSOG1* gene was constructed as follows. A 0.9 kb artificially synthesized DNA fragment (SOG1-7D GT template) that contained the *OsSOG1* coding region containing the 4th intron, 5th exon (with S306D/S316D/S322D/S338D/S340D/S398D mutations and silent mutation producing an artificial *PstI* site, cttcag to ctgcag), 5th intron, 6th exon (with S403D mutation), and 3’ UTR region was cloned into the *PacI* site in pZH_OsU6sgRNA_SpCas-wt (Endo et al., 2019) using an In-Fusion HD cloning kit (Takara). Two pairs of 20-nt annealed oligonucleotides targeting exon 4 of the *OsSOG1* gene were cloned into the *BbsI* site of the pU6gRNA-oligo (pU6::sgOsSOG1-2 and pU6::sgOsSOG1-3) (Mikami et al., 2015), respectively. These sgRNA expression cassettes (OsU6::sgOsSOG1-2 and OsU6::sgOsSOG1-3) were amplified with the primer sets listed in **Table S7**, combined in tandem and integrated into the *Asc*I site in pZH_OsU6sgRNA_SpCas-wt harboring SOG1-7D GT template by an In-Fusion reaction (Takara).

To introduce mutations in the SOG1 binding motif in the promoter region of *OsRad51A2* and *OsDMC1* via CRISPR/Cas9-mediated targeted mutagenesis, the sgRNA expression vector targeting the SOG1 binding motif on the *OsRad51A2* and *OsDMC1A* promoters were constructed as detailed above. These expression cassettes (OsU6::sgOsRad51A2 pro and OsU6::sgOsDMC1A pro) were digested with *AscI/PacI* and each ligated into pZH_OsU6sgRNA_MMCas9 (Mikami et al., 2015).

Binary vectors were transformed into *Agrobacterium tumefaciens* EHA105 (Hood et al., 1993) by electroporation (*Escherichia coli* Pulser, Bio-Rad).

### Generation of Psog1-GUS, Sog1-7A, Sog1-7D and Psgl-GUS mutant rice via GT

The GT procedure with positive-negative selection using hpt as a positive selection marker and DT-A as a negative selection marker and subsequent marker excision via *piggyBac* transposition was performed as described previously (Nishizawa-Yokoi et al., 2015). Four-week-old rice calli transformed with *Agrobacterium* harboring the GT vector were selected on callus induction medium (N6D) containing 50 mg/l hygromycin B and 25 mg/l meropenem (Wako Pure Chemical Industries) for 4 weeks. Hygromycin-resistant calli were transferred to N6D medium with hygromycin and meropenem and cultured for a further 2 weeks. Genomic DNA was extracted from hygromycin-resistant calli using Agencourt chloropure (Beckman Coulter) according to the manufacturer’s protocol and subjected to PCR analysis with KOD FX neo or KOD one (TOYOBO) using primer sets 2 and 3 (**Table S7**) shown in **Figures 1**, **2** and **S2**. The PCR amplicon from primer set 2 or 3 was sequenced using the primers listed in Table S1 to confirm the junction sequence between genomic sequence and the GUS gene or positive selection marker and the amino acid substitutions in SOG1-7A mutant introduced via GT. GT callus lines that produced PCR fragments with both primer sets 2 and 3 were transferred to regeneration medium (ReIII) containing 25 mg/l meropenem without antibiotics. Shoots regenerated from callus were transferred to Murashige and Skoog (MS) medium (Murashige and Skoog, 1962) without phytohormones. GT plants carrying the insertion of the positive selection marker in the *OsSOG1* or *OsSGL* gene were identified by PCR analysis and grown in a greenhouse. T_1_ progeny plants with homozygous insertion of the positive selection marker via GT in the *OsSOG1* or *OsSGL* gene were obtained from self-pollinating T_0_ plants, and T_2_ plants obtained from T_1_ plants were subjected to further analysis.

For marker elimination in SOG1-7A mutant calli, a portion of GT callus was transferred to N6D medium without hygromycin and meropenem and cultured for 2 weeks, before being transformed with *Agrobacterium* harboring the hyPBase expression vector pPN/hyPBase (Nishizawa-Yokoi et al., 2014). Calli transformed with the hyPBase expression cassette were selected on N6D medium containing 35 mg/l geneticin (Nacalai tesque) and 25 mg/l meropenem. Geneticin-resistant callus lines were transferred to ReIII medium containing 25 mg/l meropenem to regenerate shoots from calli. To determine whether the positive selection marker was excised by hyPBase expression from the SOG1 locus with GT-mediated amino acid mutations at 7 SQ motifs, CAPS analysis combining PCR amplification and restriction digestion was performed with genomic DNA extracted from regenerated plants. PCR fragments amplified with primer set 1 (**Figure 2A**) were digested with the *HpaI* remaining in the 5th intron of *OsSOG1* after *piggyBac* excision and sequenced with the primers shown in **Table S7** to check the presence of the desired mutations by GT and the absence of a footprint at *piggyBac* excision sites in the *OsSOG1* gene. To detect re-integration of the *piggyBac* transposon, PCR analysis was also conducted using hpt-specific primers (**Table S7**). Marker-free GT plants harboring *Hpa*I-digested PCR fragments but not hpt-specific PCR fragments were grown in a greenhouse to obtain T_1_ progenies. T_1_ plants with homozygous point mutations via GT and segregated out hyPBase expression cassette were selected by PCR, and T_2_ progenies obtained from self-pollinating T_1_ plants were subjected to further analysis.

To generate SOG1-7D mutant rice by CRISPR/Cas9-mediated GT, a GT vector harboring CRISPR/Cas9, OsU6::sgSOG1-2/3, hpt, and a SOG1-7D GT template were introduced into 4-week-old rice calli by *Agrobacterium*-mediated transformation. Following a 4-week hygromycin selection, hygromycin-resistant callus lines were subjected to DNA extraction using a one-step DNA extraction protocol (Kasajima et al., 2004). PCR fragments amplified with primers shown in **Figure S4** (**Table S7**) were digested with the *PstI* site introduced into 5th exon of *OsSOG1* gene via GT.

### Generation of mutant rice plants by the CRISPR/Cas9 system

Four-week-old rice calli transformed with *Agrobacterium* harboring a CRISPR/Cas9 expression vector were selected for 2 weeks on N6D medium containing 50 mg/l hygromycin B and 25 mg/l meropenem. Hygromycin-resistant calli were transferred to N6D medium with hygromycin and meropenem and cultured for a further 2 weeks. Genomic DNA was extracted from hygromycin-resistant calli using a one-step DNA extraction protocol (Kasajima et al., 2004).

CRISPR/Cas9-mediated mutations in the *OsSOG1* and *OsSGL* genes were detected in transgenic calli by CAPS analysis. The target region on the OsSOG1 or OsSGL gene was amplified using the primers shown in **Table S7**. PCR products were digested with *PmeI* and *Ava*II for the detection of mutations in *OsSOG1* and *OsSGL* genes, respectively, and then analyzed by agarose gel electrophoresis. Targeted mutations via CRISPR/Cas9 at SOG1 binding motifs in *OsRad51A2* and *OsDMC1A* were analyzed in transgenic calli by HMA. The target region on the *OsRad51A2* or *OsDMC1A* promoter was amplified using the primers shown in **Table S7**. PCR products were analyzed using MCE-202 MultiNA with a DNA-500 kit (Shimadzu).

Transgenic calli carrying targeted mutations via CRISPR/Cas9 at high frequency were transferred to ReIII medium containing 25 mg/l meropenem. Regenerated plants were subjected to sequencing analysis using the primers listed in **Table S7** to examine the mutation pattern at the target site. Regarding the *sog1sgl* double mutant, T_1_ plants obtained from self-pollinating T_0_ plants were subjected to PCR analysis using hpt-specific primers (**Table S7**) to select double mutant plants in which the CRISPR/Cas9 vector had segregated out. T_2_ progenies obtained from self-pollinating CRISPR/Cas9 cassette-free T_1_ plants were subjected to further analysis.

### Southern blot analysis

Genomic DNA was extracted from GT plants using the Nucleon Phytopure Extraction Kit (Cytiva) according to the manufacturer’s protocol. Genomic DNA (2 μg) was digested with *Xba*I (for Psog1-GUS mutant), *Hin*dIII (for Psgl-GUS mutant), or *XhoI* (SOG1-7A mutant) and fractionated in a 0.7% (w/v) agarose gel. Southern-blot analysis was performed according to the Digoxigenin Application Manual (Roche Diagnostics). Specific DNA probes for hpt and *OsSOG1* were synthesized with a PCR Digoxigenin Probe Synthesis Kit (Roche Diagnostics) using the primers shown in **Table S7**.

### Histochemical analysis of GUS gene expression

Psog1-GUS and Psgl-GUS mutant plants were grown at 28°C under 16 h light:8 h dark on 1/2 MS medium (pH 5.8) for 1 week. Seedlings were transferred to 1/2 MS liquid medium with or without 10 μM bleomycin and cultured for 8 h at 28°C under normal light conditions. Histochemical analysis of GUS activities in Psog1-GUS and Psgl-GUS mutant plants was performed by incubating whole seedlings in 5-bromo-4-chloro-3-indolyl glucuronide (X-gluc) buffer [100 mM sodium phosphate buffer, pH 7.0, 0.5 mM K_3_Fe(CN)_6_, 0.5 mM K_4_Fe(CN)_6_, 0.3% Triton X-100, 20% methanol, 1.9 mM X-gluc] at 37 °C overnight. The seedlings were washed with 70% ethanol until they were cleared of chlorophyll, then observed using Leica MZFIII microscope and DFC310 FX Digital color camera (Leica).

### DNA damage and oxidative stress susceptibility test

For the DNA damage susceptibility test, wild-type, Psog1-GUS, SOG1-7A, Psgl-GUS, *sog1sgl* mutant seeds were sown on 1/2 MS medium containing bleomycin at various concentrations and were cultured for 1 week, and their sensitivity evaluated by measuring root length. For the oxidative stress susceptibility test, wild-type, Psog1-GUS, SOG1-7A and sog1sgl mutant seeds were sown on 1/2 MS medium with or without 0.05 μM MV and cultured for 1-week.

### RNA extraction and microarray analysis

One-week-old wild-type, Psog1-GUS, SOG1-7A, Psgl-GUS, *sog1sgl* mutants were transferred to 1/2 MS liquid medium with or without 10 μM bleomycin and were cultured for 8 h at 28°C under normal light conditions. The shoots and roots were harvested separately and frozen immediately in liquid nitrogen. Total RNA was isolated from roots treated with or without bleomycin using an RNeasy Plant Mini Kit (Qiagen), labeled using a Low Input Quick-Amp Labeling Kit (Agilent Technologies), and hybridized to a rice 4 × 44 K custom oligo DNA microarray (Agilent Technologies) according to the manufacturer’s instructions. Hybridization microarray slides were scanned with a DNA microarray scanner G2505C (Agilent Technologies). The images generated were analyzed using Feature Extraction software version 11.5.1.1 (Agilent Technologies) applying standard normalization procedures.

### RT-PCR and qRT-PCR

Total RNA isolated from shoots and roots treated with or without bleomycin using an RNeasy Plant Mini Kit (Qiagen) was treated with DNase I (Qiagen) to eliminate any contaminating DNA. First-strand cDNA was synthesized using ReverTra Ace (TOYOBO) with an oligo (dT)_20_ primer. RT-PCR was performed with PrimeSTAR GXL DNA polymerase (Takara) using gene-specific primers (**Table S7**). Quantitative RT-PCR was conducted with a Power SYBR Green PCR Master Mix (Thermo Fisher) and Applied Biosystems ViiA 7 Real-Time PCR System (Thermo Fisher) and using gene-specific primers (**Table S8**).

### Search for SOG1 binding motif in the promoter region of DDR genes

We extracted the 1 kb regions upstream of the translation start site of genes and searched for the possible SOG1 promoter motif “CAA(N)_7_TTG” that was reported as a binding motif of Arabidopsis SOG1 using a Perl script for pattern matching.

## Author contributions

AN-Y and ST designed the research, AN-Y, AM, KI conducted the experiments, RM performed microarray analysis. TT conducted in silico analysis, AN-Y wrote the manuscript, ST edited the manuscript.

## Acknowledgments

We are deeply grateful to Dr. K. Yoshiyama (Tohoku Univ.) for insightful discussions and comments on the manuscript. We would like to thank Ms. K. Amagai, Ms. C. Furusawa, Ms. A. Nagashii, Ms. A. Sugai and Ms. R. Takahashi for experimental technical support. We are grateful to Drs. K. Uchino and H. Sezutsu (NARO) for providing hyPBase and *piggyBac* IVR vectors, and Drs. H. Saika, M. Endo, S. Hirose, H. Ichikawa (NARO), and Dr. T. Maruta (Shimane University) for helpful discussions and comments, and Dr. H. Rothnie for English editing.

This research was also supported in part by JST, PRESTO (Grant No. JPMJPR16QA), a Research Grant from NARO Gender Equality Program, Cross-ministerial Strategic Innovation Promotion Program (SIP), “Technologies for Smart Bio-industry and Agriculture” (funding agency: Bio-oriented Technology Research Advancement Institution), Cabinet Office, Government of Japan, Moonshot Research and Development Program for Agriculture, Forestry and Fisheries (JPJ009237, funding agency: Bio-oriented Technology Research Advancement Institution), and JSPS KAKENHI (Grant No.JP19K21185 and JP21K05535).

